# The circadian clock regulates *KCNH2* (hERG) promoter activity through daily temperature rhythms

**DOI:** 10.64898/2026.09.28.751769

**Authors:** Ezekiel Rozmus, Isabel G Stumpf, David Schneider, Alexander Alimov, Taylor H. Pierce, Tooraj Mirshahi, Brian P Delisle, Elizabeth A Schroder

**Author notes:** Co-corresponding senior authors: Elizabeth A. Schroder Department of Internal Medicine 1095 Veterans Drive, HSRB 262, Lexington, KY 40536-0305 Brian P. Delisle 741 S Limestone Street BBSRB B353, Lexington, KY, 40536.

## Abstract

**Background:** *KCNH2* encodes Kv11.1 channel proteins that conduct the rapidly activating delayed-rectifier K+ current (I_Kr_), which is critical for cardiac repolarization. *KCNH2* encodes two functional isoforms, Kv11.1a and Kv11.1b, via alternative transcription start sites. Kv11.1a is the principal determinant of cardiac I_Kr_ and ventricular repolarization. The circadian clock, a transcriptional-translational feedback loop that cycles with a period of ∼24 hours and drives the circadian expression of many genes, including *Kcnh2* in the mouse heart. Because daily body temperature rhythms provide a systemic signal that synchronizes cardiac circadian clocks, we tested whether physiological temperature cycles drive the circadian promoter activity of the cloned human *KCNH2* (h*KCNH2*) promoter.

**Hypothesis:** h*KCNH2* is a direct transcriptional target of the circadian clock, with temperature driving its promoter activity through BMAL1:CLOCK acting at a conserved tandem E-box.

**Methods:** We cloned the conserved proximal promoter of *KCNH2* (-1631 bp upstream of Kv11.1a exon 1) to generate h*KCNH2* promoter luciferase reporter constructs. Constructs were transfected into C2C12 myotubes and synchronized by serum shock (static 37°C) or temperature cycling (36.5–38.5°C). Bioluminescence was recorded and assessed for period, phase, and amplitude. BMAL1:CLOCK dependence was tested via dominant-negative CLOCKΔ19 co-expression.

**Results:** Temperature cycling did not exhibit the rapid damping characteristic of serum-shock-synchronized oscillations, consistent with continuous entrainment by an external zeitgeber rather than a free-running oscillator. Deletion analysis identified a conserved tandem E-box required for oscillation under both serum shock and temperature cycling, and for BMAL1:CLOCK-dependent transactivation (1.75 ± 0.21 vs. 0.86 ± 0.06 RLU, p = 0.0038). CLOCKΔ19 reduced h*KCNH2* promoter amplitude under temperature cycling without altering period.

**Conclusion:** The circadian clock regulates *KCNH2* promoter activity through daily temperature rhythms.

## Introduction

*KCNH2* encodes the pore-forming subunit of the Kv11.1 K^+^ channel proteins that conduct the rapidly activating delayed-rectifier current (I_Kr_) in the heart ^1^. I_Kr_ is a primary driver of repolarization, and reductions in I_Kr_ due to drug block or loss-of-function mutations in *KCNH2* can cause long QT syndrome (LQTS) ^1^.

In mice, cardiac *Kcnh2* expression is regulated by the circadian clock, a transcriptional-translational feedback loop in which the BMAL1:CLOCK heterodimer binds E-box elements (CANNTG) to drive expression of core clock genes including *Per1/2* and *Cry1/2*, which in turn suppress BMAL1:CLOCK activity to complete the ∼24-hour cycle ^2,3^. Beyond this core timekeeping role, BMAL1:CLOCK heterodimers drive expression of clock-controlled output genes important for local cell physiology, including several cardiac ion channels^4,5^. Cardiomyocyte-specific deletion of *Bmal1* in mice disrupts circadian expression of multiple ion channel mRNA transcripts, including *Kcnh2*, and prolongs the QT interval at slow heart rates, providing direct evidence that circadian expression of ion channel transcription contributes to normal ventricular repolarization ^4,6^.

The circadian rhythm in core body temperature (approximately 1–2°C over the 24-hour cycle) serves as a physiological time cue that entrains the phase of peripheral tissue circadian clocks including the heart ^7–9^. These temperature oscillations may represent a direct mechanistic link between daily changes in core body temperature and circadian clock regulation of cardiac ion channel promoter activity. We tested whether 24-hour temperature rhythms regulate the circadian expression of the human *KCNH2* promoter via the circadian clock mechanism.

We addressed this question by cloning the human *KCNH2* promoter (h*KCNH2*) and monitoring its promoter activity in real time using a promoter-reporter luciferase assay in synchronized C2C12 myotubes exposed to either serum shock at static culture temperature (37°C) or oscillating temperature conditions (36.5-38.5°C) mimicking physiological core body temperature rhythms. We demonstrate that oscillating temperature markedly increased h*KCNH2* circadian amplitude and drove circadian oscillations in transcription. Using systematic deletion analysis and dominant-negative clock protein expression, we identified a conserved tandem E-box element at −960 bp upstream of the Kv11.1a channel subunit translation start site as essential for both temperature-driven and circadian regulation of h*KCNH2* expression. Disruption of BMAL1:CLOCK function reduced oscillation amplitude and altered circadian period under serum shock but not under oscillating temperature, confirming that the clock machinery drives the rhythmic component of h*KCNH2* transcription.

## Materials and Methods

### Plasmid Constructs

The conserved 1631 bp human *KCNH2* 5′-promoter sequence (upstream from the Kv11.1a translation start site) was PCR-amplified from human genomic DNA and cloned into the pGL3 basic vector (Promega; 212936) to generate the full-length h*KCNH2* promoter-reporter construct (h*KCNH2*:LUC). *KCNH*2 expresses two functional isoforms, Kv11.1a and Kv11.11b^1^. The Kv11.1b isoform is generated by an alternate exon 1 and start site. The cloned 1631 bp fragment corresponds to the conserved proximal promoter of the Kv11.1a isoform. We could not identify a conserved region upstream of the Kv11.1b isoform. The sequence of the h*KCNH2* promoter-reporter construct (h*KCNH2*:LUC) was verified by DNA sequencing (ACGT DNA Sequencing Services). The tandem E-box deletion construct (Δ h*KCNH2*:LUC (−960)) was generated by site-directed mutagenesis of the full-length h*KCNH2*-luc construct using the QuikChange Site-Directed Mutagenesis Kit (Agilent; 200519) according to the manufacturer’s instructions. The tandem E-box is located approximately 960 bp upstream of the transcription start site of NM_000238.4, based on alignment of the cloned promoter fragment to the NCBI RefSeqGene record for *KCNH2* (NG_008916.1, LRG_288). All mutant constructs were verified by DNA sequencing prior to use. *Bmal1*:luc, *Per1*:luc and the CLOCKΔ19 expression plasmid^10,11^ were a kind gift from the laboratory of Dr. Karyn Esser at the University of Florida.

### Cell Culture

iCell Cardiomyocytes (Cellular Dynamics International/FUJIFILM) were thawed and maintained according to the manufacturer’s User’s Guide and cultured for 7 days prior to transfection. On the day of transfection, spent medium was aspirated and replaced with fresh iCell Cardiomyocytes Maintenance Medium at 90% of the total culture volume, and cells were incubated for 2–4 hours (37°C, 5% CO_₂_). *Bmal1*:luc and *Per1*:luc reporter plasmids were complexed with ViaFect Transfection Reagent (Promega; E4981) in Opti-MEM Reduced Serum Media (Life Technologies; 31985-062) at a 2:1 reagent (µl):DNA (µg) ratio to generate a 10X transfection complex solution, which was added to the center of each lumicycle dish containing cardiomyocytes in Maintenance Medium. Plates were incubated overnight (37°C, 5% CO_₂_), after which 100% of the medium was replaced with fresh Maintenance Medium. Cells were prepared for LumiCycle bioluminescence recording 24 hours after transfection (see cell culture section).

Mouse C2C12 myoblasts (ATCC; CRL-1772) grown in DMEM (Gibco; 11995-065) growth media (10% fetal bovine serum (Atlanta Biologicals; S11150); 1% Penicillin Streptomycin (ThermoFisher; 15140-122)) were transiently transfected with WT or mutant h*KCNH2* promoter-reporter plasmids using standard protocols for Polyjet Transfection Reagent (SignaGen; SL 100688). Transfected cells were grown in 35 mm dishes to confluence. Upon reaching confluence, cells were differentiated for 5 days in DMEM media containing 1% Penicillin Streptomycin and 2% Horse Serum (Sigma; H1138). Cells were synchronized with 50% horse serum shock or temperature oscillation^9,12^. Media was then changed to LumiCycle media containing phenol red free DMEM (Sigma; D2902); Glucose 19.4 mM (Sigma; G7528); NaHCO_3_ 4.17 mM (Sigma; S6297); HEPES Buffer 10mM (Sigma; H4034); 1% Penicillin Streptomycin; 2% Horse Serum; and 0.1 mM d-Luciferin (Biosynth; L-8220). At least two separate transfections were performed for each plasmid construct.

### Temperature Oscillation

Cells were placed in the LumiCycle with temperature oscillating between 36.5°C and 38.5°C over a 24-hour period, mimicking the physiological range of circadian variation in core body temperature. Temperature was cycled using a Nearpow timer (model T329C) with multiple programmed on/off intervals controlling power to the entire incubator, allowing the incubator to be actively heated and then passively cooled at intervals adjusted to prevent excessive cooling overnight. Because the incubator’s water jacket provides substantial thermal mass, passive cooling proceeds gradually and is influenced by ambient room temperature, whereas active heating is not. Actual incubator temperature was independently recorded throughout each experiment using a HOBO pendant data logger (Onset).

### Assessment of Circadian Promoter Activity

Circadian clocks were synchronized across cells using one of two methods: serum shock^13,14^ or temperature cycling. For serum shock synchronization, cells were treated with 50% horse serum for 2 hours, after which media was replaced with LumiCycle media and bioluminescence recording began immediately. For temperature synchronization, cells were placed in the LumiCycle with temperature oscillating between 36.5°C and 38.5°C over a 24-hour period, mimicking the physiological range of circadian variation in core body temperature ^15^. Bioluminescence was measured at 10-min intervals for 7–10 days using the LumiCycle™ (Actimetrics). Baseline-subtracted data were analyzed for circadian characteristics including period and amplitude using LumiCycle Analysis software. The period length of luminescence for each reporter gene was calculated based on the distances between peaks or troughs once synchronized (∼3–5 days). Peak phase for each replicate was determined. For temperature-synchronized cells, phase was measured relative to the trough of the temperature cycle. For serum-shocked cells, the time of the first peak occurring after 24 hours in the LumiCycle was identified and used as the start point for a 5-day analysis window.

### Dual Luciferase Assay

To assess BMAL1:CLOCK transactivation of the h*KCNH2* promoter, C2C12 myoblasts were transiently co-transfected with either the full-length h*KCNH2*:luc or Δh*KCNH2*:LUC (−960) construct, a Renilla luciferase (pRL-null) control vector (Promega; E2271) for normalization, and expression plasmids encoding BMAL1 and either wild-type CLOCK or the dominant negative CLOCKΔ19 using Polyjet Transfection Reagent (SignaGen; SL 100688). Cells were harvested 48 hours post-transfection and dual luciferase activity was measured using the Dual-Glo Luciferase Assay System (Promega; E2920). Firefly luciferase activity was normalized to Renilla luciferase activity to control for transfection efficiency. Results are expressed as relative luciferase activity.

### Temperature Monitoring

Temperature within the LumiCycle was monitored using a HOBO pendant data logger (Onset). The data logger was placed inside the LumiCycle at the beginning of each experiment and recorded temperature at 30-minute intervals throughout the measurement period. Upon completion of the experiment, the data logger was removed, and temperature data were downloaded for analysis.

### Analysis

Bioluminescence data were analyzed for circadian parameters including period and amplitude using LumiCycle Analysis software (Actimetrics). Baseline subtraction was performed using the polynomial fitting function built into the LumiCycle Analysis software. The data was subsequently smoothed with a 2-hour moving average. Circadian rhythmicity was independently verified using the JTK_Cycle algorithm, a nonparametric statistical method for detecting and characterizing rhythmic patterns in biological data ^16^. Results from both analyses were consistent for period and amplitude determinations; LumiCycle-derived parameters for these measures are reported throughout.

### Cross-correlation Analysis

For temperature-synchronized replicates, each detrended bioluminescence trace was cross-correlated against its own experimentally recorded temperature trace across a range of ±12 hours to determine peak correlation coefficient (r) and lag. Because temperature was actively heated but passively cooled, with cooling rate dependent on ambient room temperature, the temperature cycle did not consistently peak or trough at a fixed interval across experiments. The onset of the temperature rise (identified as the point of maximum positive slope in the recorded temperature trace) was more reproducible across transfections than the temperature peak or trough and was therefore used as the reference point for lag, calculated as the interval between this onset and the peak bioluminescence of each replicate. Peak correlation coefficient (r), reported separately from lag, was derived from the full-waveform cross-correlation described above and reflects overall fidelity of tracking to the temperature cycle rather than a specific timing reference.

### Analysis of publicly available ChIP-seq data

Human ChIP-seq peak calls for ARNTL/BMAL1 and CLOCK were obtained from ReMap 2022 (all-peak MACS2 sets, hg38) ^17^. BMAL1 peaks at this locus derived from GSE134972 ^18^, in which BMAL1 ChIP-seq was performed in patient-derived glioma stem cell and normal neural stem cell lines; the peaks overlapping the *KCNH2* promoter came from two independent glioma stem cell lines (GSC_387 and GSC_3565). CLOCK peaks derived from GSE96659, in which CLOCK ChIP-seq was performed on Brodmann areas 10 and 40 of adult human neocortex (five replicates per region from the same five individuals) ^19^. The tandem E-box was assigned to chr7:150,979,256–150,979,275 (GRCh38) by alignment of the cloned 1631 bp fragment to the reference genome. Peaks were considered to occupy the element only if the reported summit coordinate fell within it, since peak widths (∼150–700 bp) substantially exceed the 20 bp element.

### Statistics

All results are reported as mean ± SEM. For bioluminescence data, n is reported as the number of dishes/number of independent transfections (e.g., n = 9/3 indicates 9 dishes across 3 independent transfections). Comparisons between two groups were made using an unpaired Student’s t-test. For comparisons among three or more groups, one-way ANOVA with Tukey’s post-hoc test was used. A p value < 0.05 was considered statistically significant. All statistical analyses were performed using GraphPad Prism (version 10). Phase and coherence across replicates were analyzed using circular statistics. For each construct, individual replicate phases were converted to angles on a 24-hour clock, and the mean phase and vector length (r, a measure of phase coherence ranging from 0 for uniformly scattered phases to 1 for perfectly aligned phases) were calculated using standard circular mean methods. Circular standard deviation was calculated as √(−2 ln r) × (24/2π).

## Results

C2C12 myotubes possess a stable circadian clock and respond reliably to serum shock ^13,14^. We demonstrate here that they also respond consistently to temperature entrainment (**Supplementary Figure 1A-D**). C2C12 cells were used as the primary experimental model for these circadian studies, as hiPSC-CMs did not consistently exhibit the expected antiphase relationship between core clock promoter reporters *Bmal1*:LUC and *Per1*:LUC in our hands (**Supplementary Figure 2**). C2C12 cells further offer the advantage of endogenously expressing the Kv11.1a isoform, with greater abundance at the post-mitotic myotube stage ^20,21^, consistent with *Kcnh2* being natively expressed and regulated in this system. Use of a h*KCHN2* promoter reporter therefore assesses promoter activity of a gene already expressed in this cell type.

We compared circadian bioluminescence between the full-length h*KCNH2* promoter and a deletion construct lacking a conserved tandem E-box (Δh*KCNH2*:LUC) under serum shock synchronization. Mean baseline-subtracted bioluminescence traces recorded over 5 days are shown in **Figure 1A**, with the full-length promoter exhibiting more sustained oscillations than the Δh*KCNH2*:LUC construct. Circadian period was stable across constructs (**Figure 1B**), while oscillation amplitude was dramatically reduced with the E-box deletion (**Figure 1C**). Phase was not significantly altered by the deletion (**Figure 1D**) (h*KCNH2*:LUC n = 13/4 separate transfections; Δh*KCNH2*:LUC n = 15/4 separate transfections).

**Figure 1.**
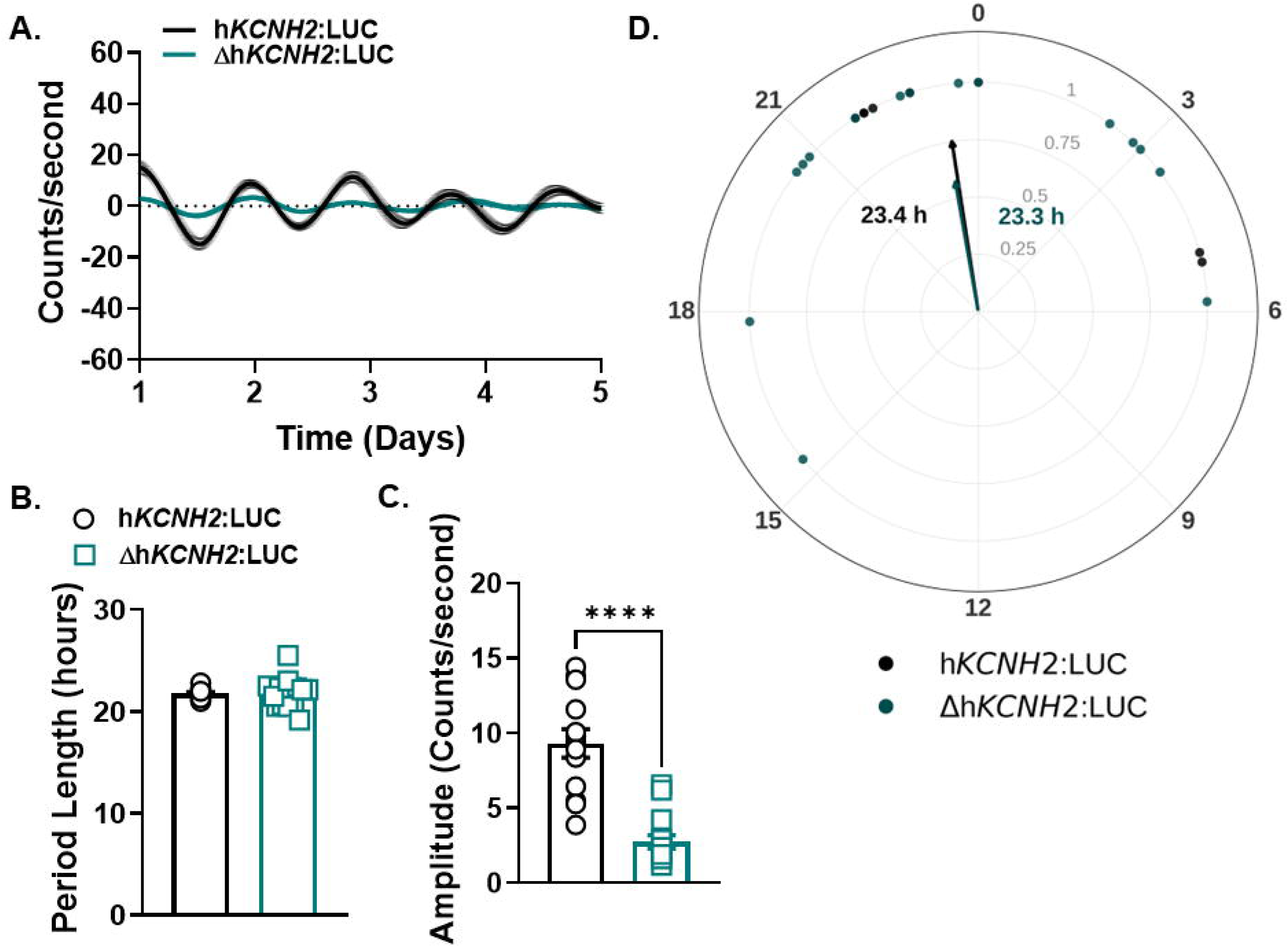
h*KCNH2*:LUC promoter activity under serum shock synchronization. **(A)** Shown are mean ± SEM baseline-subtracted bioluminescence traces for the full-length h*KCNH2*:LUC promoter and the Δh*KCNH2*:LUC deletion construct (lacking the conserved tandem E-box) recorded over 5 days following serum shock. **(B)** Circadian period length for each construct, determined by LumiCycle Analysis software (Actimetrics) is shown. **(C)** Circadian oscillation amplitude for each construct is shown. Amplitude was dramatically reduced by deletion of the tandem E-box. **(D)** Polar plot of peak phase and phase coherence across replicates (radius; vector length = mean coherence r) for each construct is shown. h*KCNH2*:LUC n = 13/4 separate transfections; Δh*KCNH2*:LUC n = 15/4 separate transfections. ****p < 0.0001.

In order to identify the specific regulatory elements responsible for circadian control of the h*KCNH2* promoter, we asked whether the tandem E-box was required for BMAL1:CLOCK-dependent transactivation. We performed dual luciferase assays in C2C12 myotubes co-transfected with either the full-length h*KCNH2*:LUC or Δh*KCNH2*:LUC construct, along with BMAL1 and either wild-type CLOCK or the dominant negative CLOCKΔ19 (**Figure 2**).

**Figure 2.**
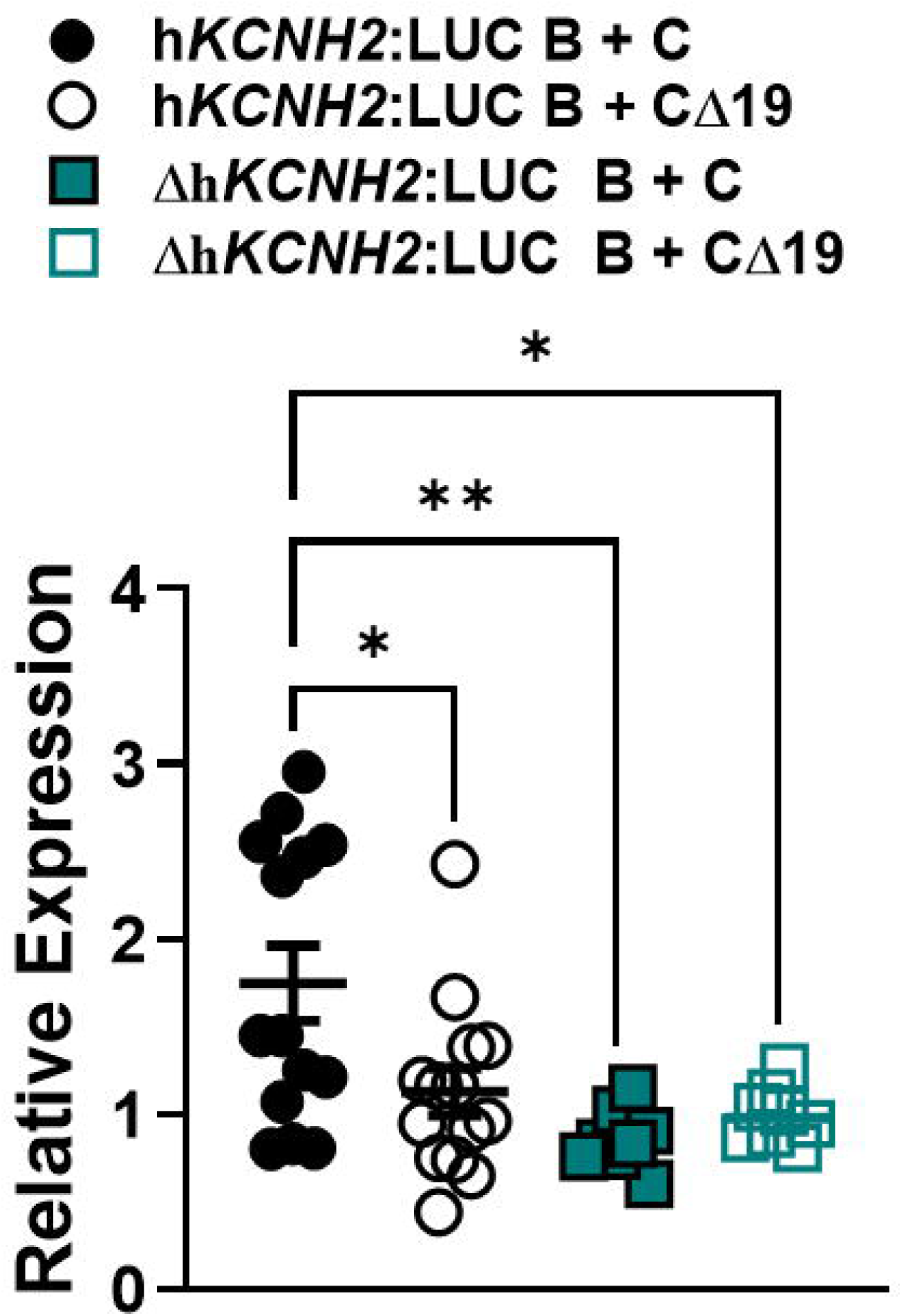
BMAL1:CLOCK-dependent transactivation of the h*KCNH2* promoter through the conserved tandem E-box. Shown are the results from the dual luciferase reporter assay in C2C12 myotubes co-transfected with either the full-length h*KCNH2*:LUC or Δh*KCNH2*:LUC (E-box deletion) construct, together with BMAL1 and either wild-type CLOCK or the dominant-negative CLOCKΔ19. Relative luciferase activity (normalized to Renilla control) is shown for each condition. BMAL1:CLOCK transactivated the full-length h*KCNH2* promoter. This transactivation was significantly reduced by co-expression of CLOCKΔ19 and by deletion of the tandem E-box. Co-expression of CLOCKΔ19 with the Δh*KCNH2*:LUC construct did not further reduce promoter activity relative to Δh*KCNH2*:LUC alone, indicating that BMAL1:CLOCK drives h*KCNH2* transcription primarily through the tandem E-box. n = 14/3 separate transfections for h*KCNH2*:LUC conditions; n = 8/3 separate transfections for Δh*KCNH2*:LUC conditions. *p < 0.05, **p < 0.01.

BMAL1:CLOCK transactivated the full-length h*KCNH2* promoter (1.75 ± 0.21 relative luciferase units (RLU)). This transactivation was significantly reduced by co-expression of CLOCKΔ19 (1.13 ± 0.13 RLU, p = 0.023) and by deletion of the tandem E-box (0.86 ± 0.06 RLU, p = 0.0038). Co-transfection of Δh*KCNH2*:LUC with CLOCKΔ19 did not further reduce promoter activity compared to Δh*KCNH2* alone (0.99 ± 0.05, p = 0.96), indicating that BMAL1:CLOCK drives h*KCNH2* transcription primarily through the conserved tandem E-box (n = 14 for h*KCNH2*:LUC conditions; n = 8 for Δh*KCNH2*:LUC conditions/3 separate transfections per condition).

To assess whether BMAL1 occupies this element in native chromatin, we queried publicly available human BMAL1 and CLOCK ChIP-seq peak sets against the genomic coordinates of the tandem E-box (chr7:150,979,256–150,979,275). Because the element spans only 20 bp, well below typical ChIP-seq peak width, peaks were classified by summit position rather than by boundary overlap. BMAL1 peaks were present at this element in two independent glioma stem cell lines, with one summit falling within the tandem E-box itself (chr7:150,979,270) and a second 32 bp away (chr7:150,979,223); two additional BMAL1 summits were located approximately 900 bp away within the cloned fragment. These were modest peak calls, as is typical of BMAL1 ChIP-seq relative to lineage-determining transcription factors. CLOCK peaks were present within the 1631 bp fragment but none had a summit at the tandem E-box, the nearest lying approximately 290 bp away (chr7:150,978,970).

We next wanted to determine if this BMAL1:CLOCK dependence extended to components of the core clock. We recorded *Bmal1*:LUC bioluminescence under serum shock in C2C12 myotubes co-expressing either wild-type CLOCK or CLOCKΔ19 (**Figure 3A**). *Bmal1*:LUC in the CLOCK (WT) background showed a tightly coherent circadian phase (18.8 h, r = 0.99, n = 4). *Bmal1*:LUC co-expressed with CLOCKΔ19 lost phase coherence (r = 0.04, n = 4), with individual replicate peaks scattered across the 24-hour cycle (**Figure 3D**). CLOCKΔ19 co-expression also lengthened circadian period (21.23 ± 0.16 *Bmal1*:LUC vs 27 ± 1.62 *Bmal1*:LUC + CLOCKΔ19, p = 0.0085); **Figure 3B**) and significantly reduced circadian amplitude (**Figure 3C**).

**Figure 3.**
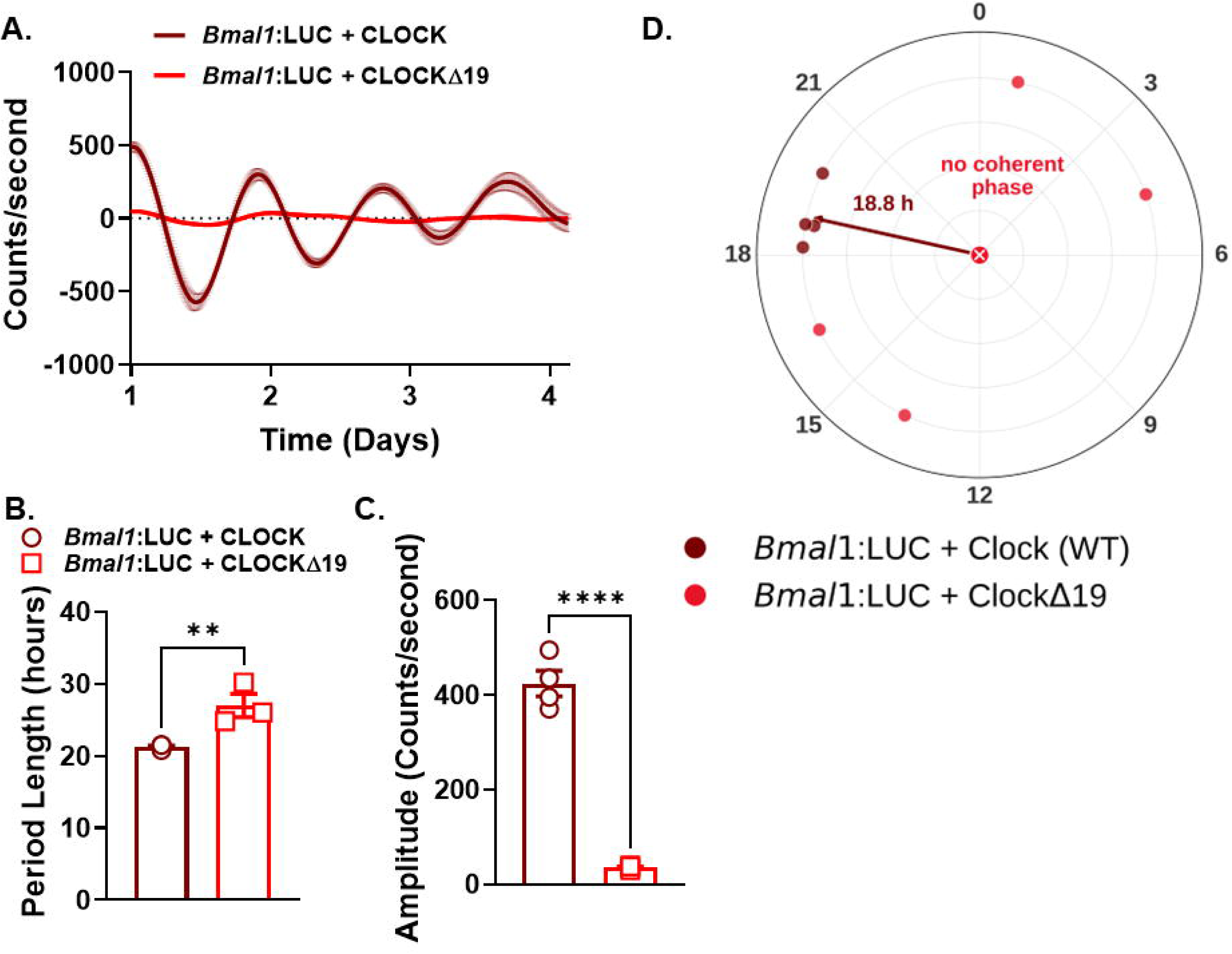
BMAL1:CLOCK dependence of core clock phase coherence under serum shock. **(A)** Shown are the mean ± SEM bioluminescence traces for *Bmal1*:LUC co-expressed with wild-type CLOCK or the dominant-negative CLOCKΔ19, recorded following serum shock. **(B)** Circadian period length is shown. CLOCKΔ19 co-expression significantly lengthened period relative to wild-type CLOCK. **(C)** Circadian oscillation amplitude is shown. CLOCKΔ19 co-expression significantly reduced amplitude relative to wild-type CLOCK. **(D)** Shown is a polar plot of peak phase and phase coherence across replicates (radius; vector length = mean coherence r). *Bmal1*:LUC with wild-type CLOCK showed a tightly coherent phase (18.8 h, r = 0.99, n = 4), while *Bmal1*:LUC co-expressed with CLOCKΔ19 showed no coherent phase (r = 0.04, n = 4), with individual replicate peaks scattered across the 24-hour cycle. n = 4 dishes per condition from a single transfection. **p < 0.01, ****p < 0.0001.

We next asked whether temperature regulates h*KCNH2* promoter activity through the same BMAL1:CLOCK-dependent mechanism identified under serum shock. Temperature was oscillated by ∼2°C over 24 hours to mimic daily changes in core body temperature. Temperature oscillation produced more reproducible oscillations in promoter activity across replicates than serum shock (peak correlation r = 0.943 ± 0.038 Serum shock vs. 0.973 ± 0.012 temperature, p = 0.039; inter-replicate lag comparable between paradigms, p = 0.623; **Supplementary Figure 3**). We compared circadian bioluminescence between the full-length h*KCNH2* promoter and the Δh*KCNH2*:LUC deletion construct under temperature cycling. Mean baseline-subtracted bioluminescence traces are shown in **Figure 4A**, with the full-length promoter exhibiting visibly higher-amplitude, more sustained oscillations than the Δh*KCNH2*:LUC construct (h*KCNH2*:LUC, 35.01 ± 1.9 vs. Δh*KCNH2*:LUC, 8.79 ± 2.3 counts/second; p < 0.0001; **Figure 4C**). Period remained consistent between constructs (**Figure 4B**), while phase was modestly but significantly delayed in the deletion construct (11.95 ± 0.76 vs. 12.40 ± 0.49 h, p = 0.027; **Figure 4D**), indicating that the tandem E-box contributes to circadian oscillation amplitude and, to a lesser extent, phase timing under temperature entrainment (h*KCNH2*:LUC n = 30/8 separate transfections; Δh*KCNH2*:LUC n = 13/3 separate transfections).

**Figure 4.**
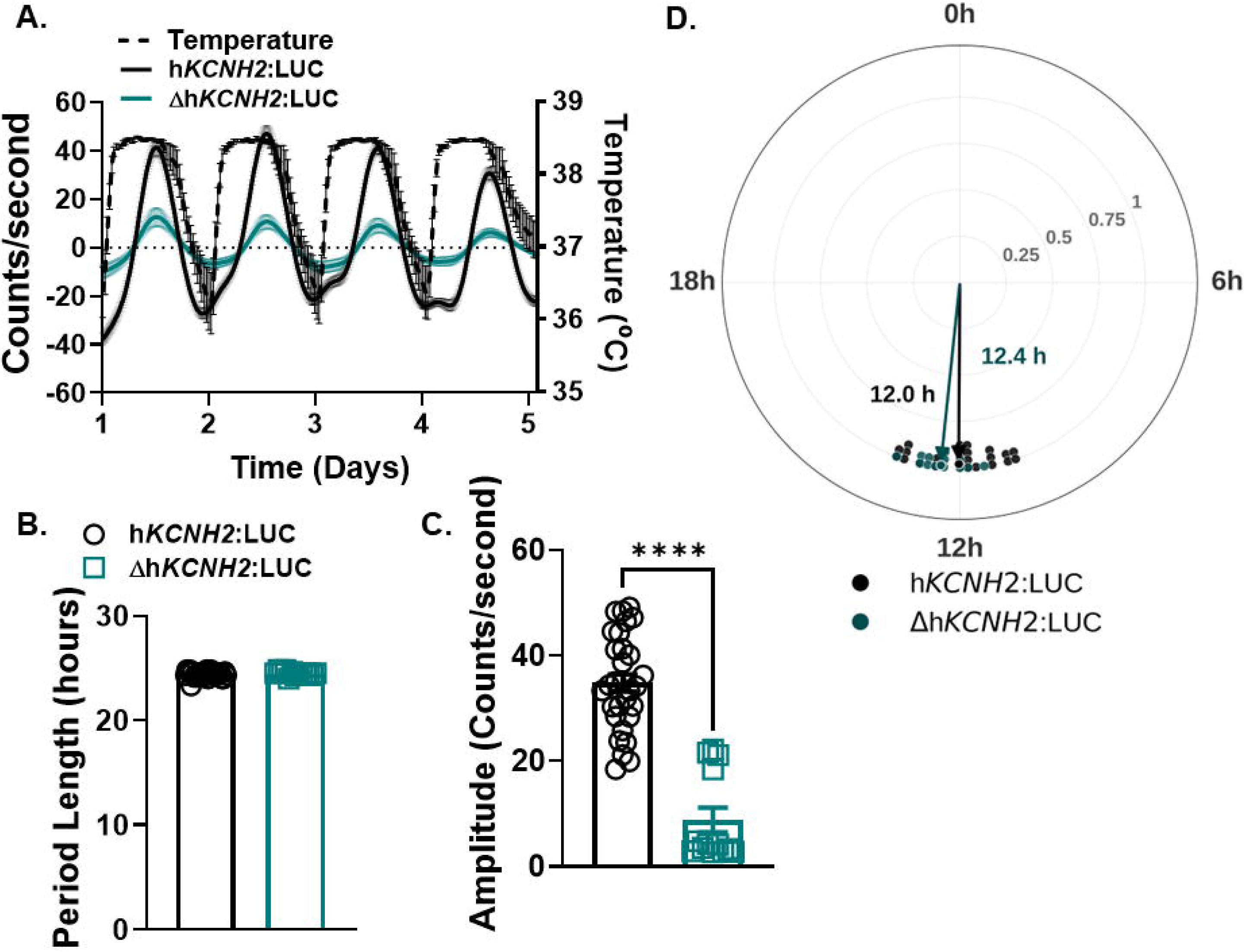
h*KCNH2* promoter activity under temperature cycling. **(A)** Mean ± SEM baseline-subtracted bioluminescence traces for the full-length h*KCNH2*:LUC promoter and the Δh*KCNH2*:LUC deletion construct, recorded over 5 days under temperature cycling (36.5– 38.5°C), shown relative to the recorded temperature trace (dashed line). **(B)** Circadian period length for each construct is shown. Period remained consistent between constructs. **(C)** Circadian oscillation amplitude for each construct is shown. Amplitude was dramatically reduced by deletion of the tandem E-box. **(D)** Polar plot of peak phase and phase coherence across replicates (radius; vector length = mean coherence r) is shown. Phase was modestly but significantly delayed in the deletion construct (11.95 ± 0.76 vs. 12.40 ± 0.49 h, p = 0.027), indicating that the tandem E-box contributes to circadian oscillation amplitude and, to a lesser extent, phase timing under temperature entrainment. h*KCNH2*:LUC n = 30/8 separate transfections; Δh*KCNH2*:LUC n = 13/3 separate transfections. ****p < 0.0001.

We next assessed entrainment to the temperature cycling stimulus by performing cross-correlation analysis between the luciferase signal from each replicate and the recorded temperature trace from its own experimental run (**Figure 5**). Cross-correlation plots are shown for the full-length h*KCNH2*:LUC construct and the Δh*KCNH2*:LUC construct (**Figure 5A**). Peak correlation coefficients were comparable between constructs (r = 0.755 ± 0.118 vs. 0.776 ± 0.079, p = 0.258; **Figure 5B**). Lag, calculated relative to the onset of the temperature rise (see Methods), was also comparable between constructs (10.86 ± 1.24 vs. 10.90 ± 0.82 h, p = 0.538; **Figure 5C**). A polar plot of peak phase (angle) and correlation strength (radius) for these same replicates confirmed that both constructs clustered tightly around a common phase with comparable correlation magnitude (**Figure 5D**).

**Figure 5.**
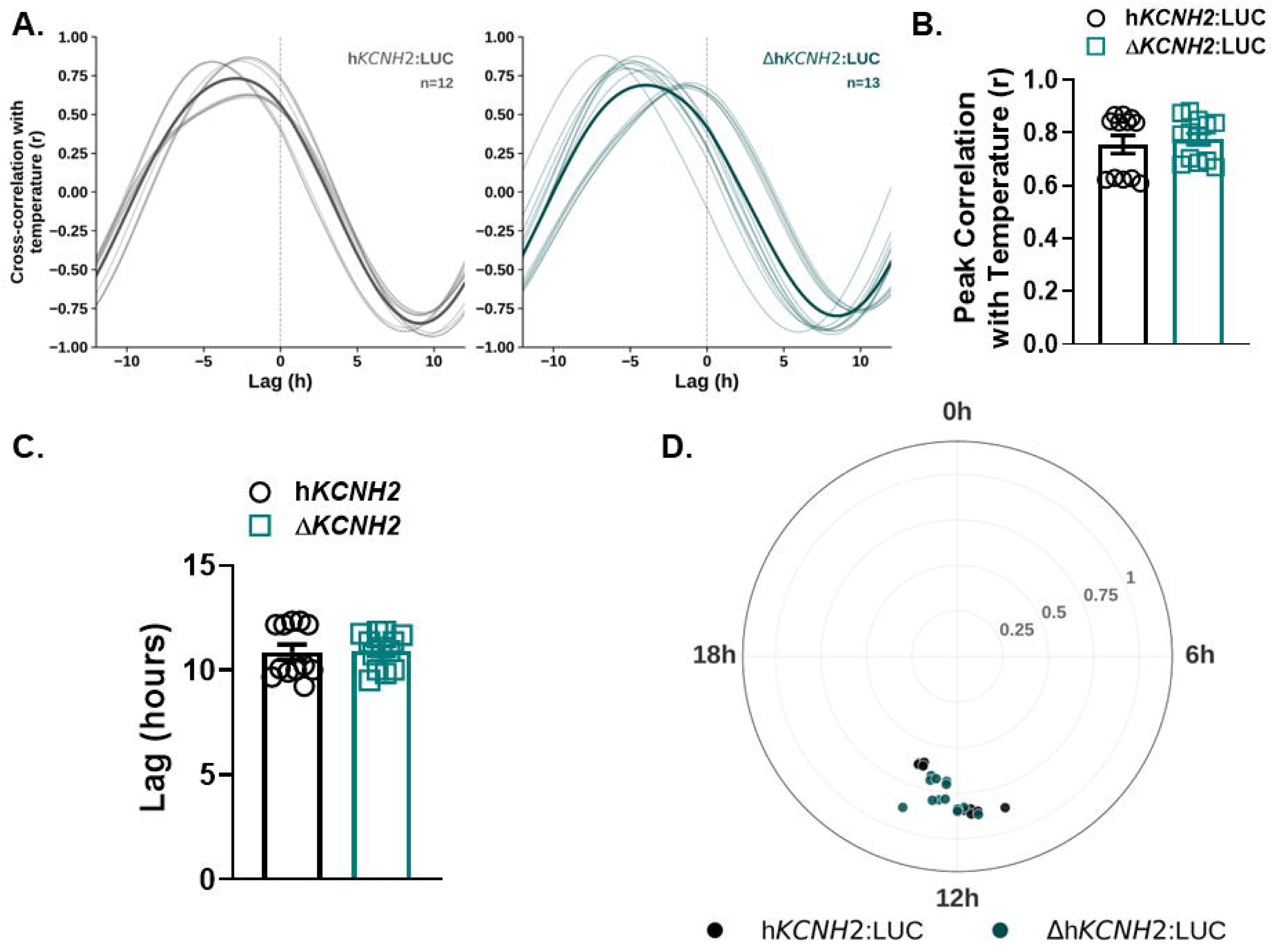
Temperature entrainment of h*KCNH2*:LUC and Δh*KCNH2*:LUC under temperature cycling. **(A)** Cross-correlation between the detrended luciferase signal for each replicate and its own recorded temperature trace, calculated across the full oscillatory waveform (lag range: −12 to +12 h) is shown. Thin lines show individual replicates; bold lines show the mean curve for h*KCNH2*:LUC (n = 12) and Δh*KCNH2*:LUC (n = 13). This analysis is shown for visualization of the overall oscillatory relationship to temperature. **(B)** Peak correlation coefficient (r) for each replicate, taken from the cross-correlation analysis is shown. **(C)** Lag is reported relative to the onset of the temperature rise rather than the full cross-correlation waveform (see Methods). Because temperature was actively heated but passively cooled, the onset of the temperature rise was more reproducible across experiments than either the temperature peak or full-waveform cross-correlation. Lag values in **(C)** are therefore not directly comparable to the lag implied by the peak position of the curves in **(A)**. **(D)** Shown is a polar plot of peak phase and peak correlation strength for replicates from the transfections in which both constructs were assayed in parallel.

Having established that deletion of the conserved tandem E-box does not significantly alter entrainment coherence or lag relative to the temperature stimulus, despite a modest phase delay detected in the intrinsic rhythm, we next asked whether BMAL1:CLOCK activity drives the circadian component of h*KCNH2* promoter oscillations in real time. We performed bioluminescence recordings in C2C12 myotubes expressing the full-length h*KCNH2*:LUC construct together with either wild-type CLOCK or ClockΔ19 under temperature cycling conditions (**Figure 6**). ClockΔ19 expression significantly reduced oscillation amplitude compared to wild-type CLOCK when co-transfected with h*KCNH2*:LUC (9.16 ± 1.98 vs. 31.31 ± 3.99 counts/second, p < 0.001, n = 11 h*KCNH2* + CLOCKΔ19, n = 7 h*KCNH2* + CLOCK, 3 separate transfections each). CLOCKΔ19 co-expression did not significantly change circadian period (24.39 ± 0.27 vs. 24.67 ± 0.14 h, p = 0.38) or phase (12.1 ± 0.6 vs. 12.6 ± 1.5 h, p = 0.37).

**Figure 6.**
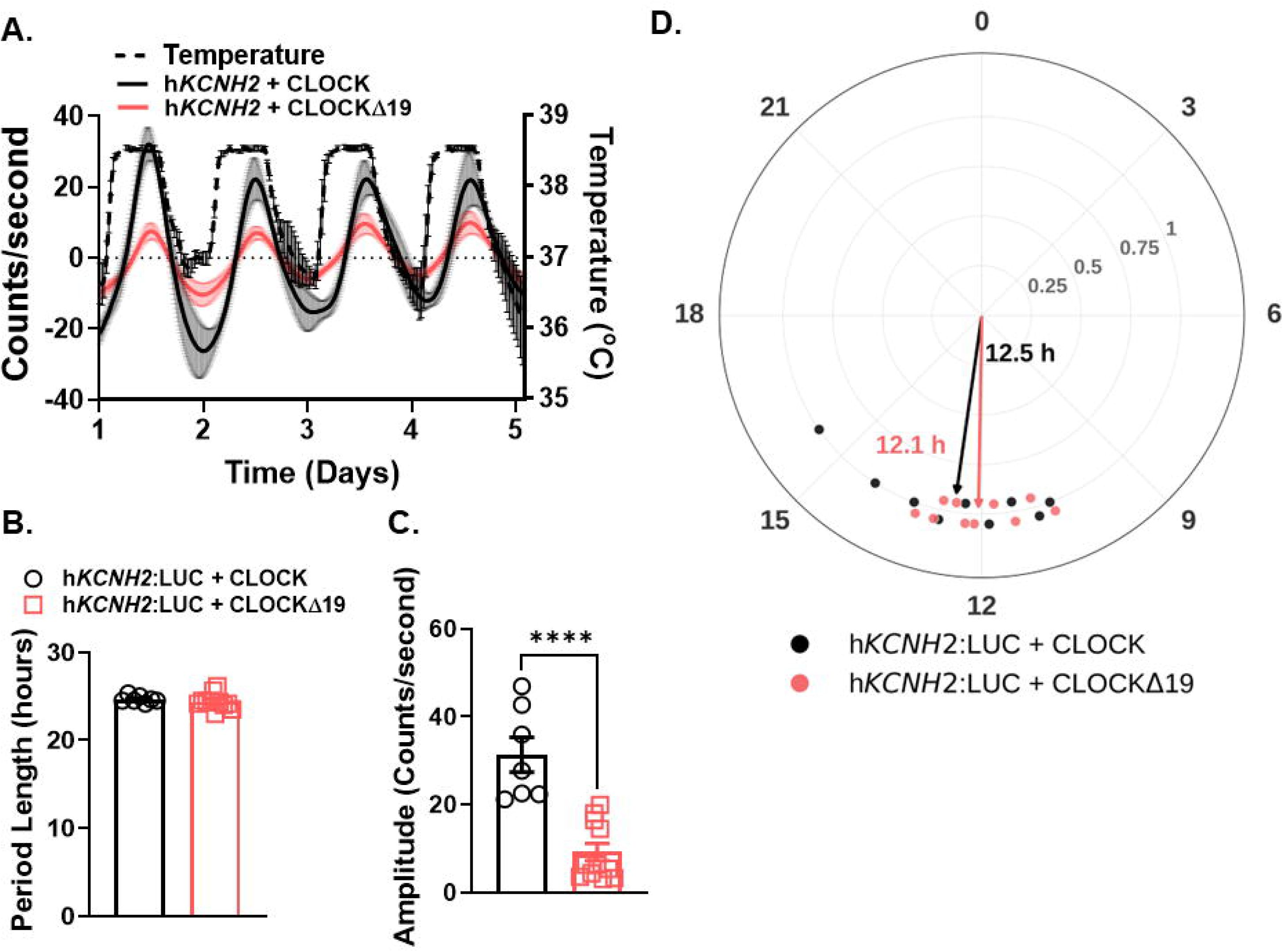
BMAL1:CLOCK drives the circadian component of h*KCNH2* promoter oscillations under temperature cycling. **(A)** Mean ± SEM bioluminescence traces for h*KCNH2*:LUC co-expressed with wild-type CLOCK or the dominant-negative CLOCKΔ19, recorded under temperature cycling, shown relative to the recorded temperature trace (dashed line). **(B)** Circadian period length is shown. CLOCKΔ19 co-expression did not significantly change period (24.39 ± 0.27 vs. 24.67 ± 0.14 h, p = 0.38). **(C)** Circadian oscillation amplitude is shown. CLOCKΔ19 co expression significantly reduced amplitude relative to wild-type CLOCK (9.16 ± 1.98 vs. 31.31 ± 3.99 counts/second, p < 0.001). **(D)** Polar plot of peak phase and phase coherence across replicates (radius; vector length = mean coherence r) is shown. Phase was not significantly altered by CLOCKΔ19 expression. n = 7 h*KCNH2*:LUC + CLOCK, n = 11 h*KCNH2*:LUC + CLOCKΔ19, 3 separate transfections each. ****p < 0.0001.

We next asked whether temperature cycling, as an external entraining signal, could sustain a coherent phase even when the internal CLOCK-dependent feedback loop is disrupted. Under temperature cycling, *Bmal1*:LUC co-expressed with WT CLOCK showed a coherent phase (19.8 h, r = 0.80, n = 9). Unlike the complete loss of phase coherence observed under serum shock (**Figure 3**), *Bmal1*:LUC co-expressed with CLOCKΔ19 under temperature cycling retained a measurable but weaker coherent phase (21.1 h, r = 0.64, n = 8; **Figure 7**) without a significant change in circadian period. This indicates that physiological temperature cycling can partially entrain circadian phase independent of an intact CLOCK-dependent feedback loop, whereas under serum shock the coherent phase/period under the same genetic disruption is lost.

**Figure 7.**
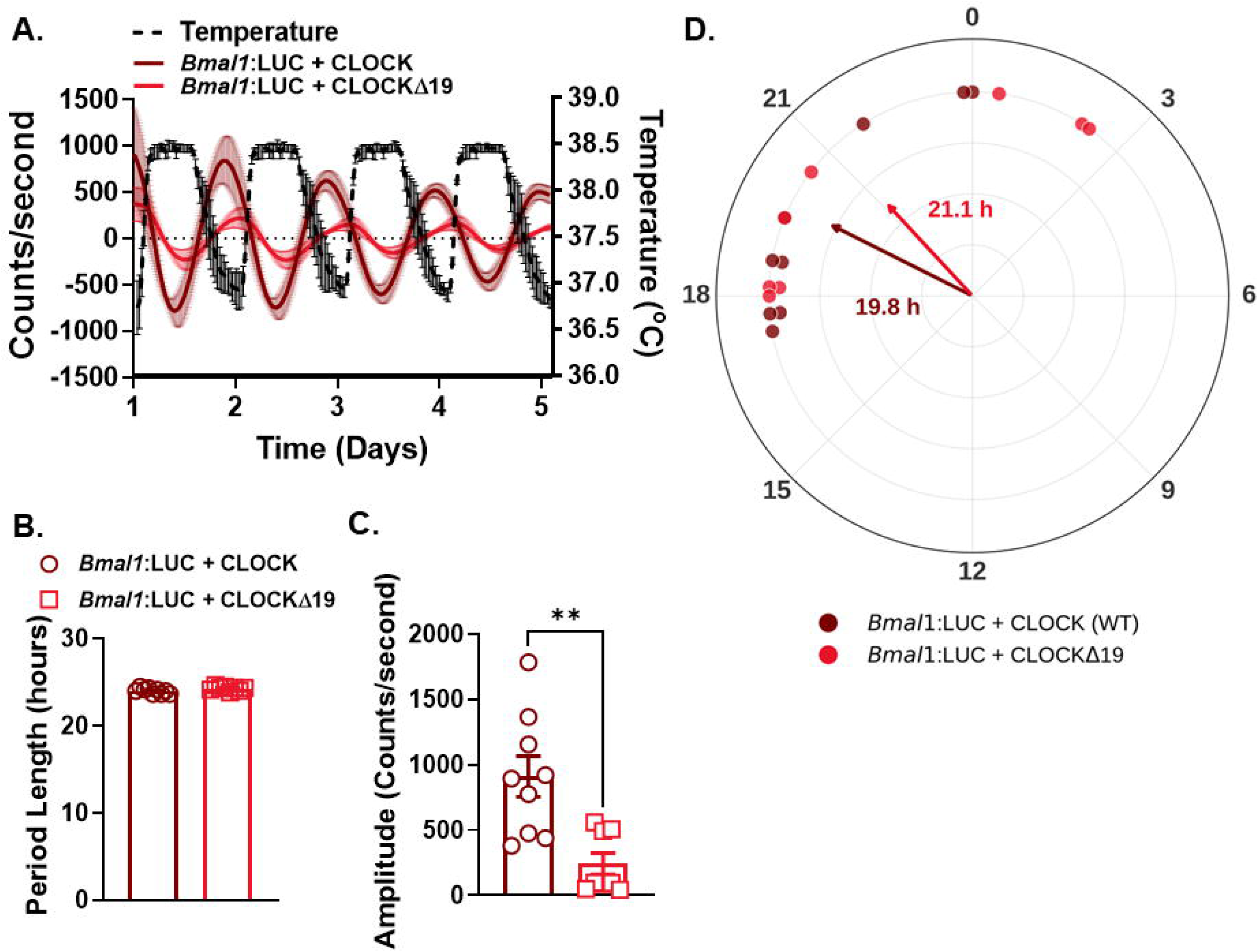
Temperature cycling partially sustains circadian phase independent of intact CLOCK function. **(A)** Mean ± SEM bioluminescence traces for *Bmal1*:LUC co-expressed with wild-type CLOCK or the dominant-negative CLOCKΔ19, recorded under temperature cycling, shown relative to the recorded temperature trace (dashed line). **(B)** Circadian period length was not different for *Bmal1*:LUC with CLOCKΔ19 co-expression. **(C)** Circadian oscillation amplitude was significantly reduced for *Bmal1*:LUC with CLOCKΔ19. **(D)** Polar plot of peak phase and phase coherence across replicates (radius; vector length = mean coherence r) is shown. *Bmal1*:LUC with wild-type CLOCK showed a coherent phase (19.8 h, r = 0.80, n = 9). In contrast to the loss of phase coherence observed under serum shock (Figure 3), *Bmal1*:LUC co-expressed with CLOCKΔ19 retained a measurable, though weaker, coherent phase under temperature cycling (21.1 h, r = 0.64, n = 8), indicating that physiological temperature cycling can partially sustain circadian phase independently of an intact CLOCK-dependent feedback loop. n = 9 *Bmal1*:LUC + CLOCK, n = 8 *Bmal1*:LUC + CLOCKΔ19/3 separate transfections. **p < 0.01.

## Discussion

The main findings of this study are that physiological temperature oscillations regulate circadian h*KCNH2* transcription through the molecular clock, and that a highly conserved tandem E-box element at −960 bp in the proximal h*KCNH2* promoter is necessary for BMAL1:CLOCK-dependent circadian transcription and the resulting increases in amplitude. Using real-time bioluminescence monitoring under serum shock and physiological temperature cycling, we demonstrate that deletion of the conserved tandem E-box dramatically reduced oscillation amplitude without altering period, establishing this element as a primary cis-regulatory determinant of circadian h*KCNH2* transcription. Disruption of BMAL1:CLOCK function by the dominant negative CLOCKΔ19 mutant reduced oscillation amplitude in cells carrying the full-length h*KCNH2* promoter under temperature cycling without altering period. Parallel experiments using a *Bmal1*:LUC reporter confirmed that CLOCKΔ19 reduced amplitude under both temperature cycling and serum shock, and lengthened period specifically under serum shock, the canonical CLOCKΔ19 phenotype (18), validating that the construct retained its expected dominant-negative activity in this system. These findings, together with the dual luciferase transactivation data, demonstrate a role for the clock machinery in driving the rhythmic component of h*KCNH2* transcription through this element.

### The tandem E-box at −960 as a primary cis-regulatory element for circadian h*KCNH2* transcription

Prior work from our laboratory demonstrated that *Kcnh2* mRNA exhibits circadian oscillations in the mouse heart and that this rhythmicity is lost in hearts with cardiomyocyte-specific deletion of *Bmal1* (iCSΔ*Bmal1*^-/-^), implicating direct clock regulation of *Kcnh2* transcription. We further showed that the *Kcnh2* promoter is transactivated by co-expression of BMAL1 and CLOCK in heterologous assays, and that the promoter drives circadian luciferase activity in real-time bioluminescence recordings ^4^. However, the specific cis-element responsible for this regulation had not been identified. The present study closes this gap by demonstrating that the conserved tandem E-box is essential for circadian h*KCNH2* promoter oscillations.

Deletion of the conserved tandem E-box reduced oscillation amplitude by 70.6% under serum shock and 74.9% under temperature cycling, while leaving period intact in both, indicating that this element is required for circadian oscillation of the h*KCNH2* promoter regardless of synchronization method, but not for the underlying timekeeping mechanism itself. To directly test whether BMAL1:CLOCK transactivates the h*KCNH2* promoter through this element, we performed dual luciferase assays using the dominant negative CLOCKΔ19 mutant, which competes with wild-type CLOCK for BMAL1 binding without driving transcription. Both CLOCKΔ19 expression and deletion of the conserved tandem E-box significantly reduced BMAL1:CLOCK-driven h*KCNH2* promoter transactivation and combining the two manipulations produced no additional reduction. This is consistent with recent structural work showing that the CLOCK exon 19 coiled-coil domain, absent in CLOCKΔ19, contributes to cooperative BMAL1:CLOCK binding at tandem E-boxes ^22^. Disrupting either the tandem element or the CLOCK domain that engages it may impair the same underlying mechanism, explaining why the two manipulations are non-additive. These data indicate that BMAL1:CLOCK transactivates the h*KCNH2* promoter primarily through the conserved tandem E-box. Consistent with this, publicly available human ChIP-seq data^17^ place a BMAL1 peak summit at the tandem E-box^18^, reproducibly across two independent human cell lines, indicating that BMAL1 can occupy this element in native chromatin. These are modest peak calls from cells in which KCNH2 is unlikely to be highly expressed, and no CLOCK summit was present at the element^19^; the available BMAL1 and CLOCK datasets derive from different studies and cell types, and no cardiac or myogenic dataset was available. This analysis therefore supports, but does not establish, direct occupancy, and confirmation of BMAL1:CLOCK binding at this element in a cardiac system will require chromatin immunoprecipitation.

The importance of tandem rather than single E-box architecture is consistent with evidence from other circadian gene promoters. Studies of core clock gene promoters including *Per1*, *Per2*, and *Per3* have shown that a direct repeat of E-box-like elements, rather than a single canonical E-box, is the minimal cis-element required for strong cell-autonomous circadian oscillation, with spacing between the two elements being critical for function ^23^. Our finding that the h*KCNH2* tandem E-box at −960 is essential for circadian expression extends this principle to a cardiac ion channel gene and suggests that tandem E-box architecture may be a conserved feature of clock-controlled genes requiring strong, tissue-specific circadian regulation.

### Temperature cycling partially sustains circadian phase independent of intact CLOCK function

A striking finding of this study is that the requirement for functional CLOCK differed by entrainment cue. Under serum shock, disruption of CLOCK function by CLOCKΔ19 completely abolished phase coherence in *Bmal1*:LUC (r = 0.04), with individual replicates scattering across the entire 24-hour cycle. However, under temperature cycling *Bmal1*:LUC co-expressed with CLOCKΔ19 retained partial, measurable phase coherence (r = 0.64), despite the same genetic disruption. This indicates that physiological temperature cycling can partially substitute for an intact CLOCK-dependent feedback loop in sustaining circadian phase, whereas in the absence of an external timing cue like in serum shock this same disruption is not tolerated.

This distinction may help reconcile our findings with in vivo models of cardiomyocyte-autonomous clock disruption. Cardiomyocyte-specific expression of the same CLOCKΔ19 mutant (CCM mice) attenuates diurnal heart rate variation and diminishes rhythmic expression of clock output genes and clock-controlled ion channels (e.g., Cav1.3) in the heart while clock function remains normal in other peripheral tissues ^24–26^. Notably, several of these cardiac phenotypes are described as attenuated rather than completely abolished. Because the heart in vivo is continuously exposed to core body temperature oscillations, an entraining signal absent in serum-shock synchronization, our finding that temperature cycling can partially sustain circadian phase despite disrupted CLOCK function offers a plausible mechanistic explanation for this partial, rather than complete, loss of rhythmicity observed in cardiac-specific clock-disruption models in vivo. This raises the possibility that core body temperature acts as a compensatory entraining cue that partially preserves clock output in cardiac tissue even when the intrinsic transcriptional feedback loop is compromised.

### Physiological temperature cycling amplifies circadian h*KCNH2* promoter activity

A major finding of this study is that oscillating temperature between 36.5–38.5°C, mimicking physiological core body temperature rhythms, markedly increases overall h*KCNH2* promoter activity compared to static culture conditions. These two effects, increased overall promoter activity and enhanced circadian amplitude, are related but distinct. The increase in overall activity may reflect a direct transcriptional response to temperature oscillations that is at least partially independent of the clock, consistent with the persistence of some promoter activity even when BMAL1:CLOCK function is disrupted. The enhanced circadian amplitude under temperature cycling, by contrast, reflects clock-dependent rhythmic transcription mediated through the conserved tandem E-box, as demonstrated by its dependence on intact BMAL1:CLOCK function and its loss by E-box deletion.

The conserved tandem E-box was essential for overall promoter activity under oscillating temperature conditions, suggesting this element serves as a convergence point for both temperature-responsive and clock-driven transcriptional inputs to the h*KCNH2* promoter. The mechanisms by which temperature directly activates transcription through this element, whether through temperature-sensitive transcription factor binding, chromatin remodeling, or indirect signaling pathways, remain to be defined.

### h*KCNH2* promoter phase corresponds to core clock timing and in vivo cardiac rhythms

To place h*KCNH2* promoter timing in the context of the core molecular clock, we compared peak phase across *Bmal1*:LUC, *Per1*:LUC, and h*KCNH2*:LUC under temperature cycling. h*KCNH2* phase-clustered closely with *Per1* (12.0 h vs. 9.6 h) rather than with *Bmal1* (21.1 h), consistent with h*KCNH2* being regulated downstream of the core negative-feedback arm of the clock (**Figure 8**). This phase relationship matched published *Kcnh2* mRNA rhythms in mouse heart in vivo ^27^, supporting the physiological relevance of the C2C12 findings and addressing a key limitation of promoter-reporter systems generally, that clock-driven transcriptional timing observed in a heterologous or immortalized cell model may not reflect the true phase relationships present in intact cardiac tissue. The concordance between our in vitro phase data and independently published in vivo transcriptomic timing ^27^ suggests that the C2C12 system despite lacking the full cardiomyocyte transcriptional and chromatin environment faithfully preserves the phase of clock-driven h*KCNH2* regulation.

**Figure 8.**
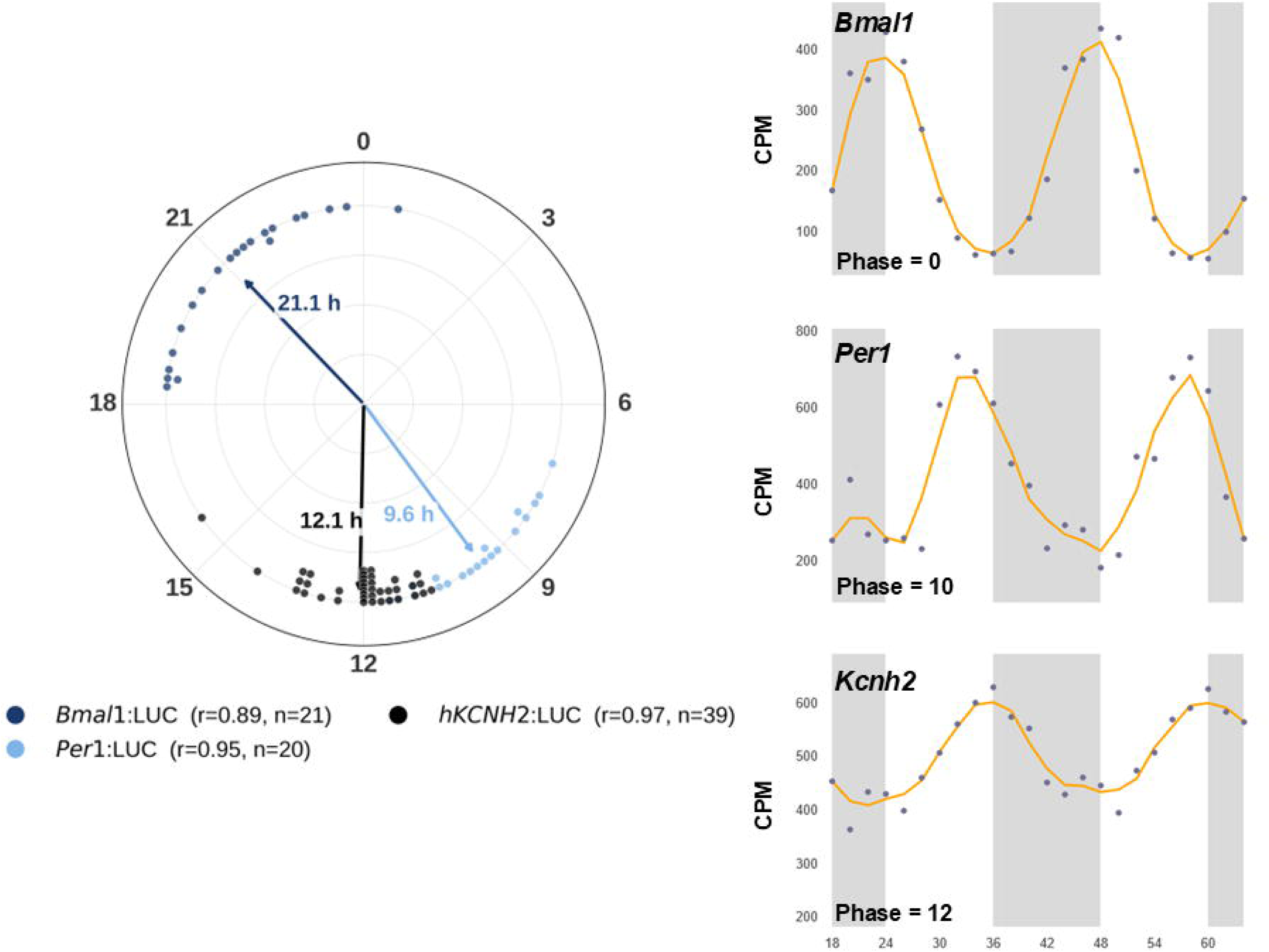
h*KCNH2* promoter phase aligns with core clock gene timing and matches in vivo circadian rhythms. Left panel. Polar plot comparing peak phase and phase coherence (radius; vector length = mean coherence r) for *Bmal1*:LUC, *Per1*:LUC, and h*KCNH2*:LUC under temperature cycling is shown. h*KCNH2* phase-clustered closely with *Per1* (12.1 h vs. 9.6 h) rather than with *Bmal1* (21.1 h), consistent with h*KCNH2* being regulated downstream of the core negative-feedback arm of the circadian clock. **Right panel.** *Bmal1*, *Per1*, and *Kcnh2* mRNA expression (counts per million, CPM) in mouse heart across circadian time (CT), obtained from CircaDB ^27^. Shaded regions indicate the dark (active) phase. Fitted phase is shown for each gene in CT hours: *Bmal1* phase = 0, *Per1* phase = 10, *Kcnh2* phase = 12. Because core body temperature is itself entrained to the rest-activity cycle, a trough occurs during the rest (light) and a peak during the active phase (dark). Phase relative to CT in vivo and phase relative to the temperature cycle in vitro (left panel) reflect similar underlying physiological timing. The relative phase relationship between these three genes parallels their C2C12 promoter-reporter counterparts, supporting the physiological relevance of the in vitro findings. *Bmal1*:LUC n = 21, *Per1*:LUC n = 20, h*KCNH2*:LUC n = 39.

### Limitations

Several limitations of this study should be noted. First, experiments were performed in C2C12 myotubes rather than primary cardiomyocytes. C2C12 cells were selected specifically because they exhibit strong temperature-entrainable circadian oscillations and are a well-validated model for circadian promoter regulation. However, they do not replicate the full complement of transcription factors and chromatin environment present in cardiomyocytes, and findings will require validation in more physiologically relevant cardiac cell models. Notably, the phase of h*KCNH2* promoter activity observed under temperature cycling is consistent with the timing of *Kcnh2* RNA expression rhythms reported in mouse heart in vivo, supporting the physiological relevance of the phase relationships identified in this system ^27–29^. Future work using hiPSC-CM cultures, in which circadian rhythmicity is more reliably established, may allow these findings to be extended to a human cardiomyocyte model.

Our ClockΔ19 experiments provide an initial dissection of clock-dependent and clock-independent contributions to *hKCNH2* temperature sensitivity. Residual oscillation amplitude persisting under CLOCKΔ19 relative to wild-type CLOCK suggests a clock-independent contribution to temperature-driven promoter activity, with the remainder attributable to CLOCK:BMAL1-dependent signaling. However, because CLOCKΔ19 may not completely eliminate residual CLOCK:BMAL1 activity, this separation is only approximate. Future studies using clock-deficient cardiomyocytes or inducible *Bmal1* deletion models would allow more precise dissection of these two regulatory components. Promoter-reporter assays measure transcriptional activity and do not directly assess I_Kr_ density or channel function. Whether the amplitude differences observed in promoter activity translate to proportional changes in I_Kr_ density in cardiomyocytes remains to be determined. The ChIP-seq data used to assess BMAL1 occupancy derive from non-cardiac, non-myogenic human cells, as no cardiac or skeletal muscle BMAL1 or CLOCK dataset was available. These data cannot establish occupancy of this element in cardiomyocytes.

### Conclusions

This study demonstrates that physiological temperature oscillations markedly increase overall h*KCNH2* promoter activity and that a conserved tandem E-box at −960 bp is the primary cis-regulatory element mediating both this temperature-driven response and BMAL1:CLOCK-dependent circadian transcription. Disruption of BMAL1:CLOCK function reduced h*KCNH2* oscillation amplitude under temperature cycling without altering period, confirming that the clock machinery drives rhythmic h*KCNH2* transcription through this element. Parallel *Bmal1*:LUC experiments confirmed the expected period-lengthening effect of the CLOCKΔ19 construct under serum shock, validating its dominant-negative activity in this system. These findings provide a molecular framework for understanding how temperature and the cardiac circadian clock converge to regulate h*KCNH2* transcription, and how genetic variation in this regulatory element may modify arrhythmia risk.

## Acknowledgements

This work was supported by National Heart Lung and Blood Institute grants R01HL141343 (Delisle) R01HL172813 (Delisle and Schroder). The content is solely the responsibility of the authors and does not necessarily represent the official views of the NIH. Claude AI (Sonnet 4.6) was used for formatting and copyediting purposes.

## Figure Legends

**Supplemental Figure 1. Validation of temperature entrainment of the core molecular clock in C2C12 myotubes. (A)** Circadian period length for *Bmal1*:LUC and *Per1*:LUC under temperature cycling are not different. **(B)** Circadian oscillation amplitude for *Bmal1*:LUC and *Per1*:LUC under temperature cycling are shown. **(C)** Polar plot of peak phase and phase coherence across replicates (radius; vector length = mean coherence r). *Bmal1*:LUC and *Per1*:LUC showed the expected near-antiphase relationship (21.1 h vs. 9.6 h; r = 0.89, n = 21 and r = 0.95, n = 20, respectively), confirming that C2C12 myotubes respond appropriately to temperature entrainment. **(D)** Mean ± SEM bioluminescence traces for *Bmal1*:LUC and *Per1*:LUC relative to the recorded temperature trace (dashed line) are shown. n = 20–21/6 separate transfections.

**Supplemental Figure 2. hiPSC-CMs do not exhibit the expected antiphase relationship between core clock reporters. (A)** Circadian period length for *Bmal1*:LUC and *Per1*:LUC in hiPSC-CMs under temperature cycling is shown. **(B)** Circadian oscillation amplitude for *Bmal1*:LUC and *Per1*:LUC in hiPSC-CMs under temperature cycling is shown. **(C)** Polar plot of peak phase and phase coherence across replicates (radius; vector length = mean coherence r) are shown. *Bmal1*:LUC and *Per1*:LUC peaked within approximately 1 hour of each other (18.4 h vs. 17.3 h, n = 3 each) rather than the ∼12-hour antiphase relationship expected of a mature molecular clock. **(D)** Mean bioluminescence traces for *Bmal1*:LUC and *Per1*:LUC relative to the recorded temperature trace (dashed line) are shown. n = 3/2 transfections.

**Supplemental Figure 3. Temperature cycling produces more reproducible oscillations than serum shock. (A)** Cross-correlation of detrended h*KCNH2*:LUC signal for each replicate against the group-average trace its own construct, shown separately for serum shock (n = 10) and temperature cycling (n = 30). Thin lines show individual replicates; bold lines show the group mean. **(B)** Peak correlation coefficient (r) with the group mean for each replicate are shown. Temperature cycling produced significantly higher peak correlation than serum shock indicating more reproducible oscillations across biological replicates. **(C)** Lag (hours) between each replicate and the group-average trace are shown. Lag was comparable between paradigms (p = 0.623, ns). n = 10 (serum shock), n = 30 (temperature)/3-8 separate transfections.

