## Supplement Figures 1-3 for "The circadian clock regulates *KCNH2* (hERG) promoter activity through daily temperature rhythms"

### Slide 1
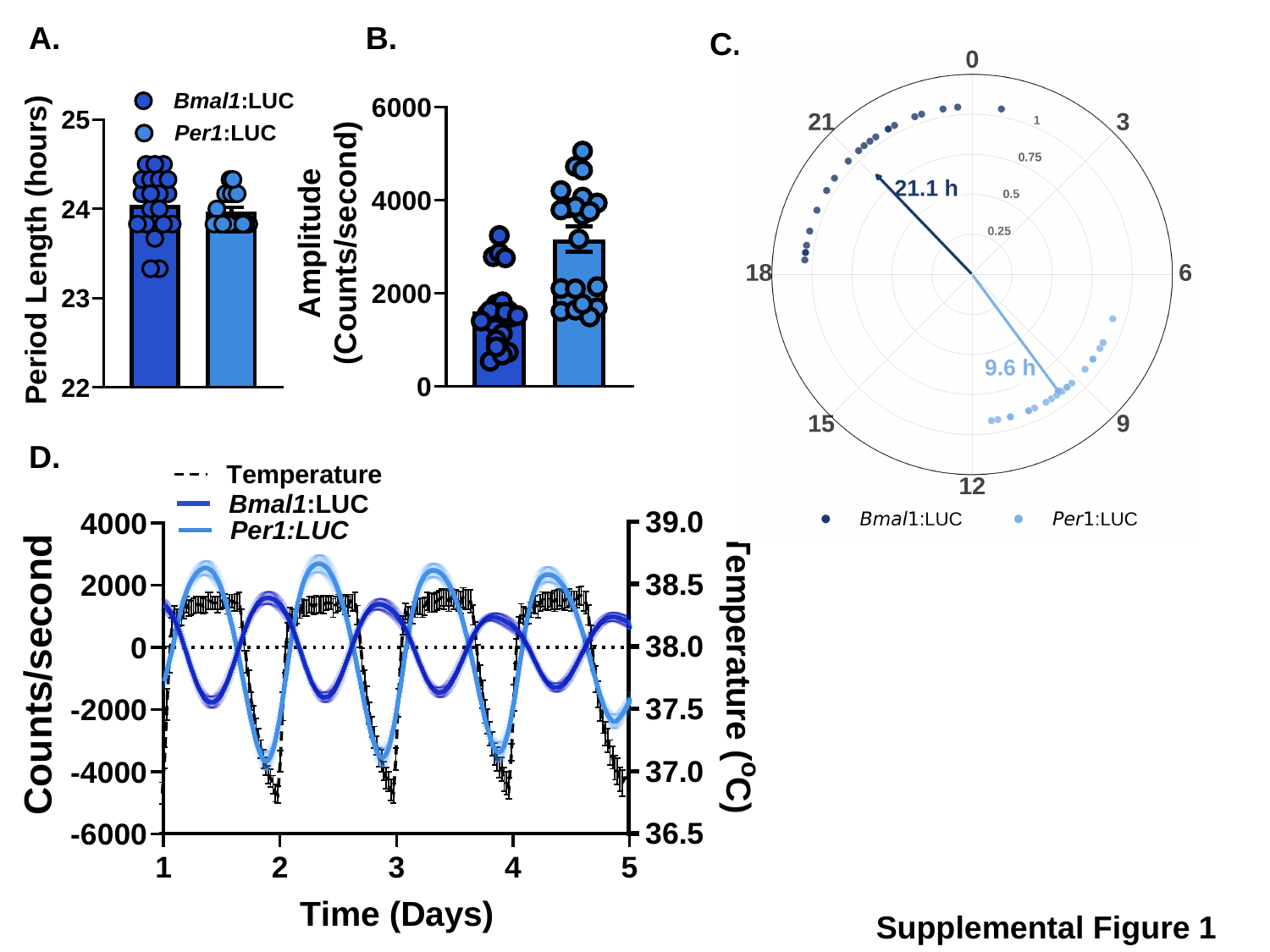

A.
B.
C.
D.
Supplemental Figure 1

### Slide 2
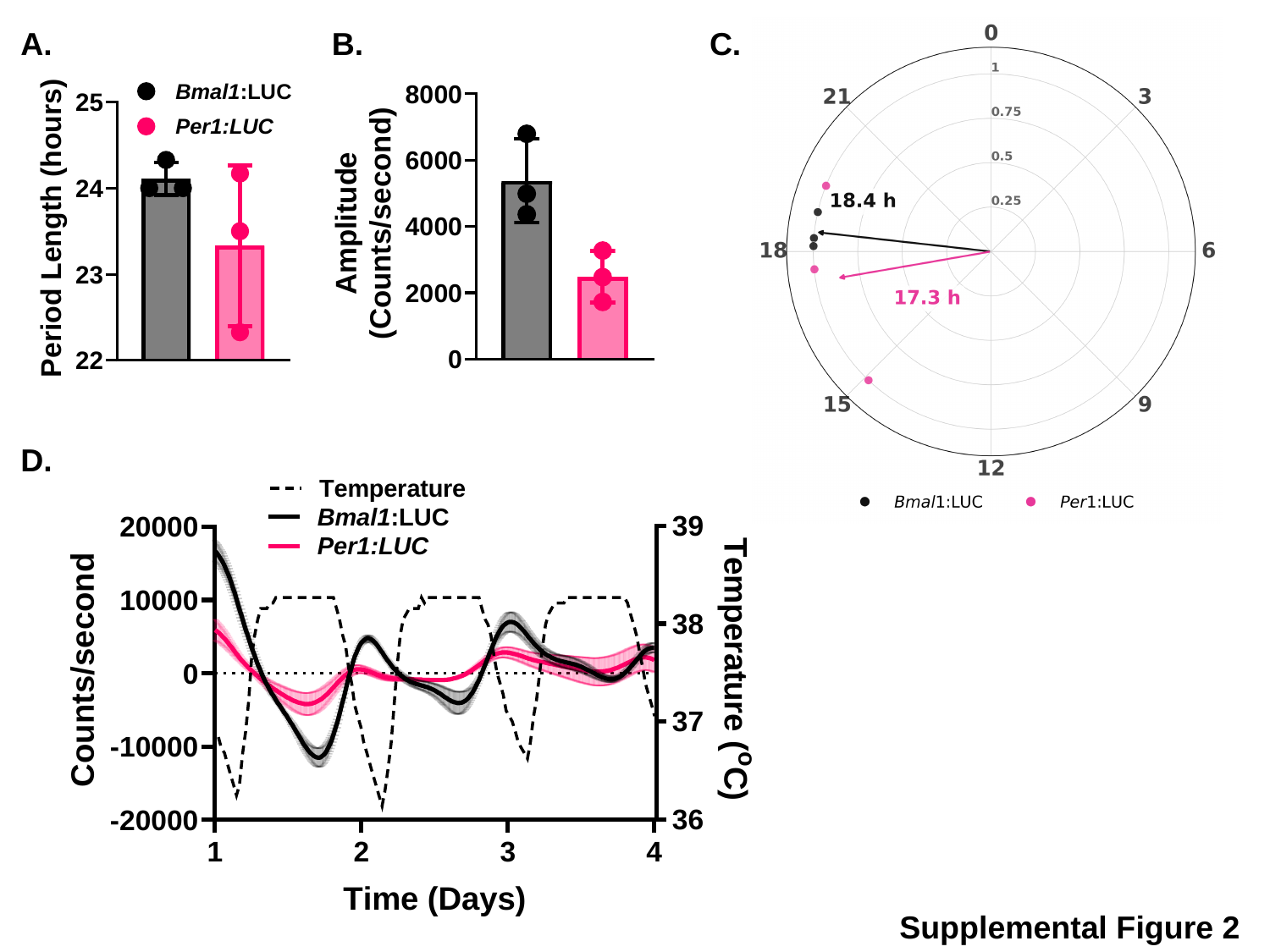

A.
B.
C.
D.
Supplemental Figure 2

### Slide 3
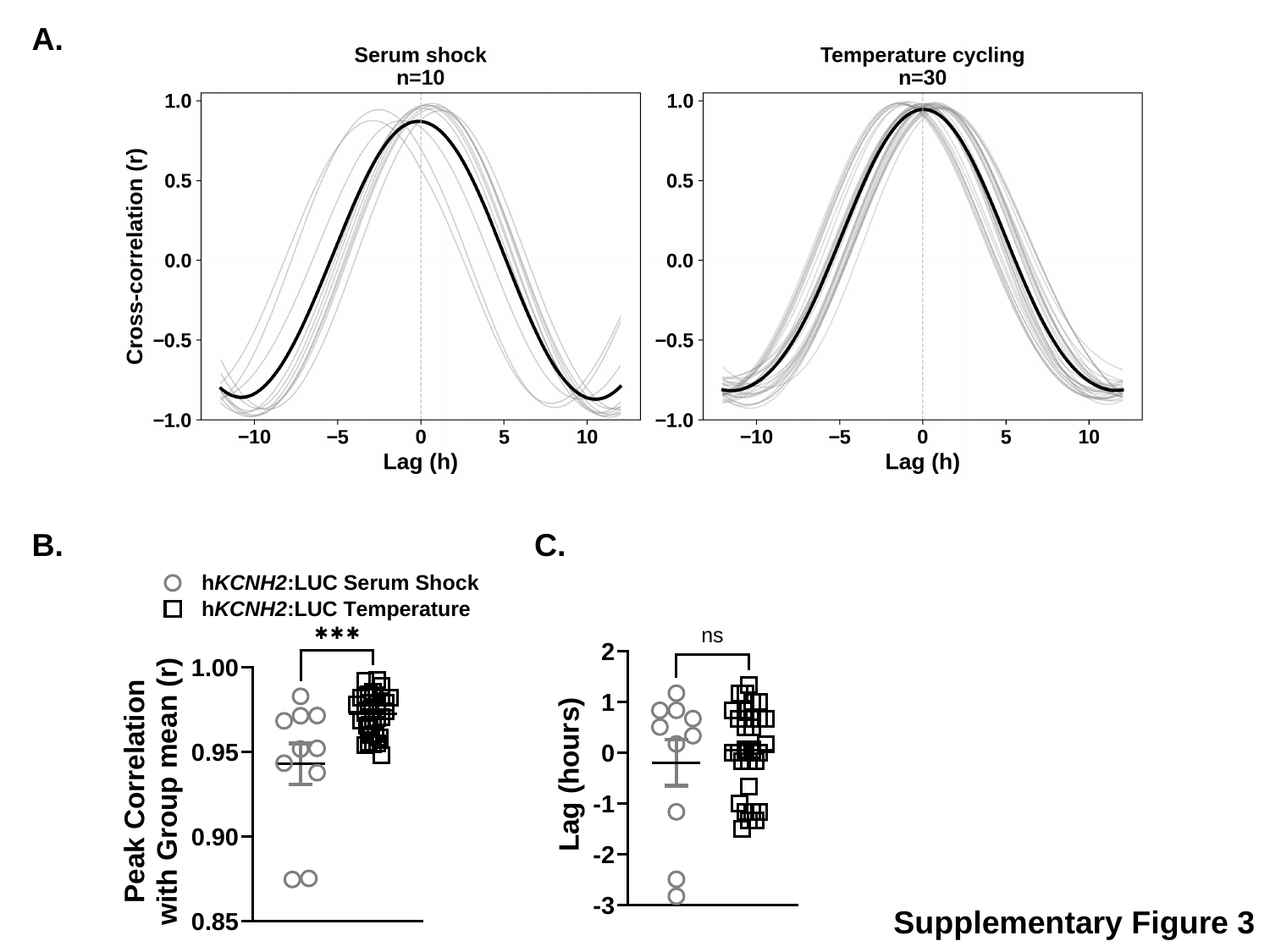

A.
B.
C.
Supplementary Figure 3
